# CellPhePy: a Python implementation of the CellPhe toolkit for automated cell phenotyping from microscopy time-lapse videos

**DOI:** 10.1101/2025.01.07.631669

**Authors:** Laura Wiggins, Stuart Lacy, Graeme Park, Joanne Marrison, Ben Powell, Beth Cimini, Peter O’Toole, Julie Wilson, William J. Brackenbury

## Abstract

We previously developed the CellPhe toolkit^1^, an open-source R package for automated cell phenotyping from ptychography time-lapse videos. To align with the growing adoption of python-based image analysis tools and to enhance interoperability with widely used software for cell segmentation and tracking, we developed a python implementation of CellPhe, named CellPhePy. CellPhePy preserves all of the core functionality of the original toolkit, including single-cell phenotypic feature extraction, time-series analysis, feature selection and cell type classification. In addition, CellPhePy introduces significant enhancements, such as an improved method for identifying features that differentiate cell populations and extended support for multiclass classification, broadening its analytical capabilities. Notably, the CellPhePy package supports CellPose segmentation and TrackMate tracking, meaning that a set of microscopy images are the only required input with segmentation, tracking and feature extraction fully automated for downstream analysis, without reliance on external applications. The workflow’s increased flexibility and modularity make it adaptable to different imaging modalities and fully customisable to address specific research questions. CellPhePy can be installed via PyPi or GitHub, and we also provide a CellPhePy GUI to aid user accessibility.

## Introduction

Recent years have seen the proliferation of open-source software for the analysis of microscopy images. These tools provide a diverse range of capabilities, ranging from segmentation and phenotypic feature extraction using CellProfiler^2^, CellPose^3^, StarDist^4^ and Ilastik^5^, to cell tracking for monitoring temporal changes to cellular phenotypes using software such as TrackMate^6^, CellTracker^7^ and DeepCellTracking^8^. With increasingly specialised analysis requirements, it is becoming more common to integrate multiple tools within a workflow, allowing researchers to take advantage of each software’s strengths for specific tasks. A key limitation of this approach is the variety of programming languages used by different software, which introduces additional complexity and necessitates advanced programming skills to integrate these tools effectively into a cohesive workflow.

Napari^9^ addresses this issue by providing a platform that supports analysis plugins, enabling tools from disparate fields to be used in combination with each other with ease. Notably, Napari is a python-based platform, which has prompted community efforts to port existing software into compatible python plugins. This trend extends beyond Napari itself, as the growing popularity of python for image analysis tasks has led to a broader movement in the community to create python versions of various tools and libraries, as well as increasing functionality to use tools in headless mode within scripts without the need for external graphical user interfaces, exemplified by tools such as Ilastik and CellPose. The integration of Docker containers further streamlines this process by allowing users to package tools along with their dependencies, ensuring consistent and reliable execution in headless environments, and tools such as BiaPy^10^ have successfully made use of this.

We previously developed the CellPhe toolkit^1^, an open-source R package for automated cell phenotyping from ptychography time-lapse videos. Recognising the current trend towards python-based analysis and the increased potential of interoperability with existing bioimage analysis tools, we sought to develop a python-based implementation of CellPhe, called CellPhePy, that retains its core functionality of single-cell phenotypic feature extraction, time-series analysis, feature selection and cell type classification. We have also made significant enhancements to the code, including a novel approach for feature selection that ensures the identification of discriminating features is more robust. Additionally, our implementation extends beyond binary classification and offers support for multi-class classification, allowing for the simultaneous characterisation and classification of multiple cell types. CellPhePy also facilitates automated CellPose segmentation and TrackMate tracking, stages that previously required manual intervention through the FIJI graphical user interface. This improvement ensures that the only necessary prerequisite for analysis is a set of microscopy time-lapse images, with all additional steps carried out automatically. CellPhePy can be downloaded and installed via PyPi or from the CellPhePy GitHub repository where you can also find a Jupyter notebook for running the CellPhePy workflow and visualising outputs. We also offer a graphical user interface (GUI) that can be easily installed through the terminal or using Docker. This GUI enables users to run the entire CellPhePy pipeline with minimal coding required, streamlining the process and making it more accessible to a broader user base.

## Results

### The CellPhePy python package

The original CellPhe R package required users to segment and track cells through external software before importing the data into the CellPhe pipeline. Although it supported TrackMate tracking tables, this process required users to be proficient with external tools and introduced additional time investment, creating a barrier to entry for using CellPhe. To address this issue, the CellPhePy package automates the process of CellPose segmentation and TrackMate tracking, meaning that a stack of time-lapse images is the only required input (**Figure 1**). As well as saving the user’s time by removing the need to manually use external software, providing a fully automated interface can save computational time too as it facilitates the deployment of CellPhe on more powerful resources such as High Performance Computing (HPC) clusters or Cloud compute. Following segmentation, the user can export their segmentation masks as well as single-cell regions of interest (ROIs) that are compatible with ImageJ’s ROI manager to visualise segmentation performance. Furthermore, CellPhePy’s tracking function allows users to select the most suitable TrackMate algorithm for their application, choosing from SimpleSparseLAP, SparseLAP, Kalman, AdvancedKalman, NearestNeighbor, or Overlap. The output file from this segmentation and tracking pipeline is then compatible with the remaining CellPhe functions within the package. These encompass calculation of phenotypic features from each cell on each frame of the time-lapse, as well as summarisation of single-cell time series to quantify temporal changes to cellular phenotypes. Cell populations can then be analysed through CellPhe’s feature selection method of “separation scores”^1^ to identify discriminatory variables, integrated XGBoost^11^ for cell type classification, and hierarchical clustering to identify heterogeneous phenotypes within a sample. Enhanced modularity allows users to customise the segmentation and tracking workflow by importing their own segmentation masks or tracking tables for use within the CellPhe pipeline. Additionally, users can integrate their own single-cell features into the code before performing time series summarisation, enabling the analysis of temporal changes in custom features. This flexibility provides users with greater control over the workflow, allowing for tailored analyses to suit their specific needs.

**Figure 1.**
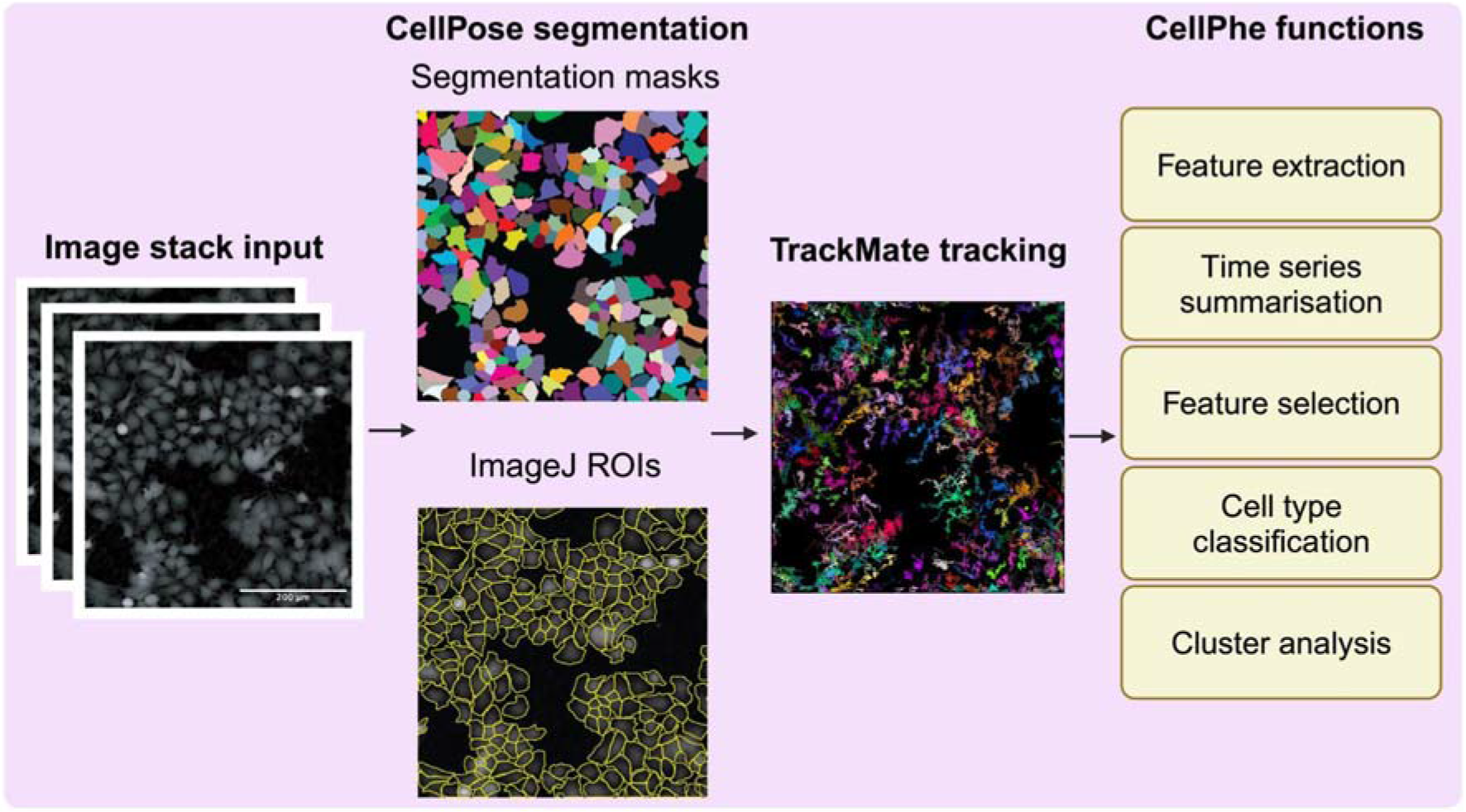
A summary of the CellPhePy package workflow. A stack of microscopy time-lapse images is the required input for the CellPhePy workflow. Each image is then automatically segmented using CellPose, where segmentation masks and ImageJ compatible ROI files can be output to assess segmentation performance. Cells are then automatically tracked through TrackMate, and tracking information fed into CellPhe’s own functions for downstream analysis.

The CellPhePy python package is available to install through PyPi at https://pypi.org/project/cellphe/ or through GitHub at https://github.com/uoy-research/CellPhePy. Note that the PyPi version 0.4.0 was used to obtain the results presented here. The GitHub page features detailed documentation in the README and a comprehensive tutorial that guides users through the workflow, from importing images to classifying phenotypic changes induced by drug treatment. In addition, a Jupyter notebook is provided, allowing users to execute the workflow with minimal coding intervention. The notebook highlights customisable parameters and offers the ability to visually inspect outputs, enabling users to explore and experiment with the results interactively.

### Improved identification of discriminatory features

The CellPhePy package includes an improved method for feature selection, enabling more reliable identification of features that effectively discriminate between different cell populations. The CellPhe pipeline determines whether a feature is discriminatory or not through calculation of a separation score that minimises within-group variance whilst maximising between-group variance, a higher separation score represents a more discriminatory feature. The original CellPhe release provided a method for determining the optimal separation threshold through classification, where the threshold was set to the lowest value that did not result in significant loss of classification accuracy. This approach involved testing classification results for discrete threshold values, which was computationally expensive and limited to fixed points. It also depended on the choice of classification model and its parameters, making it sensitive and potentially inconsistent across different setups. To address these limitations, the CellPhePy package now employs the elbow method, a proven technique for determining the optimal number of clusters in cluster analysis^12^. In this context, the elbow of the plot of ordered separation scores is identified to determine the minimum number of features that provide the greatest separation between cell populations. This approach ensures that the most discriminatory features are selected efficiently, avoiding overfitting while maximising the distinction between groups.

An example of this approach is provided in **Figure 2**, where the aim of the analysis is to identify the phenotypic features that best discriminate between strongly metastatic MDA-MB-231 cells and weakly metastatic MCF-7 cells (**Figure 2a**). A total of 1083 features are extracted through CellPhePy, and a plot of all features, coloured by feature category (size, shape, texture, movement or density) is provided in **Figure 2b**. Through the elbow method, the optimal separation threshold was determined to be 0.09, with 74 features having separation score greater than or equal to this threshold. The top scoring features were primarily related to cellular movement, characterising increased migration of MDA-MB-231 cells in comparison to MCF-7. Such features quantified temporal changes to displacement, surface area covered by cell trajectories, velocity and track length. Kernel density plots of the top-scoring features for each category are shown in **Figure 2c**, demonstrating how the separation scores correlate with the degree of overlap between the feature distributions of the two cell populations. To illustrate how the 74 retained features collectively characterise cell types, we compare PCA score plots using all 1083 features versus only the 74 highest-scoring features (**Figure 2d**). When all features are included, the plots show significant overlap between the cell populations, with no clear separation even along the first principal component. After feature selection, however, there is distinct separation between the cell populations, with greatest separation achieved along the first principal component.

**Figure 2.**
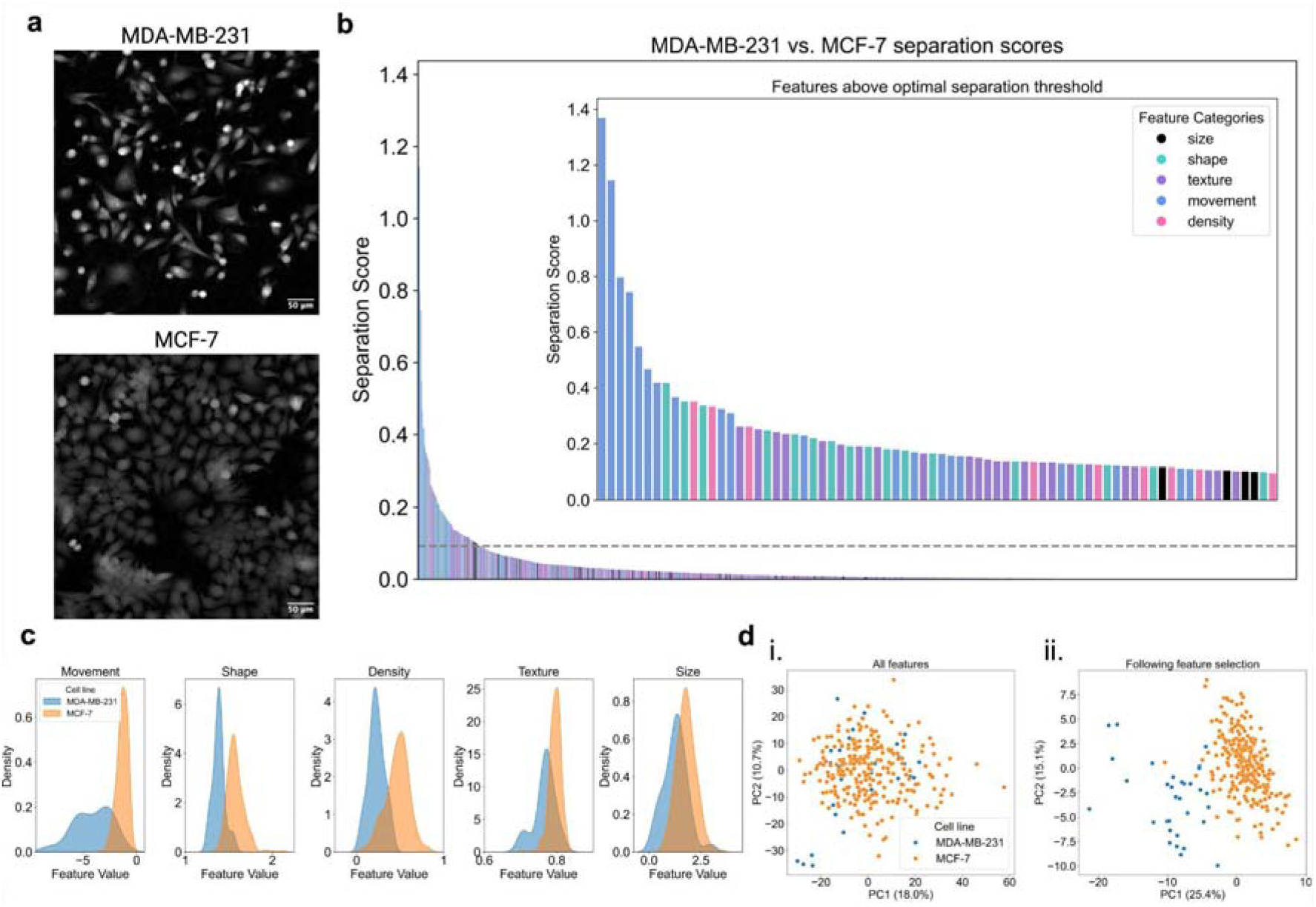
Identifying phenotypic features that characterise MDA-MB-231 and MCF-7 cells. **a**. Snapshots taken from ptychographic timelapse videos of MDA-MB-231 and MCF-7 monocultures. **b**. Barplot of separation scores for all 1083 phenotypic features extracted through CellPhe, coloured by feature category (size, shape, texture, movement, density). The grey dashed line represents the optimal separation threshold (0.09) calculated through the elbow method, with the inset plot providing a zoom in of the 74 features with separation scores greater than or equal to this threshold. **c**. Kernel density plots of the top-scoring features for each category, demonstrating relationship between separation score and separation of feature distributions. **d**. PCA scores plots when (i) all 1083 features were included and (ii) only the 74 highest-scoring features were included, showing greater separation of cell lines following feature selection.

### CellPhePy enables multiclass characterisation and classification

While the initial release of CellPhe successfully enabled the characterisation and classification of two cell types, biological experiments often require the analysis of multiple cell types simultaneously. To accommodate this, we adapted the calculation of separation scores in CellPhe to identify features that provide optimal separation across multiple cell types at once. In addition, CellPhe’s original approach to classification was through an ensemble of LDA, Random Forest and SVM classifiers, where final classification was based on majority vote, treating each classifier’s predictions equally. Whilst this works for binary classification, where one class will always receive a majority vote, it was not suitable for multiclass cases. To address this, we replaced the discrete three classifier ensemble with XGBoost, which is an ensemble of decision trees that can inherently handle multiclass problems as well as automatically weighting its members’ predictions.,

CellPhePy’s multiclass classification capabilities offer flexibility in defining and adjusting groupings to suit the specific goals of an analysis. In our example, we classify and characterise four breast cancer cell lines (MDA-MB-231, MCF-7, SkBr3, and BT-474) alongside a healthy breast epithelial cell line (MCF-10A). **Figure 3b**(i) illustrates how multiclass feature selection effectively separates these cell lines, with an optimal separation threshold of 0.45, retaining 97 phenotypic features. When the cell lines are relabelled as “cancer” for the breast cancer lines and “healthy” for MCF-10A, separation scores are recalculated to reflect disease status rather than individual cell lines. This relabelling results in an optimal separation threshold of 0.2, with 74 features identified as most discriminatory, further improving group separation in **Figure 3b**(ii). Lastly, with the healthy cell line removed, the remaining breast cancer cell lines are relabelled by their clinical molecular subtype: TNBC for MDA-MB-231 and MDA-MB-468, Luminal A for MCF-7, and Her2+ for SkBr3.

**Figure 3.**
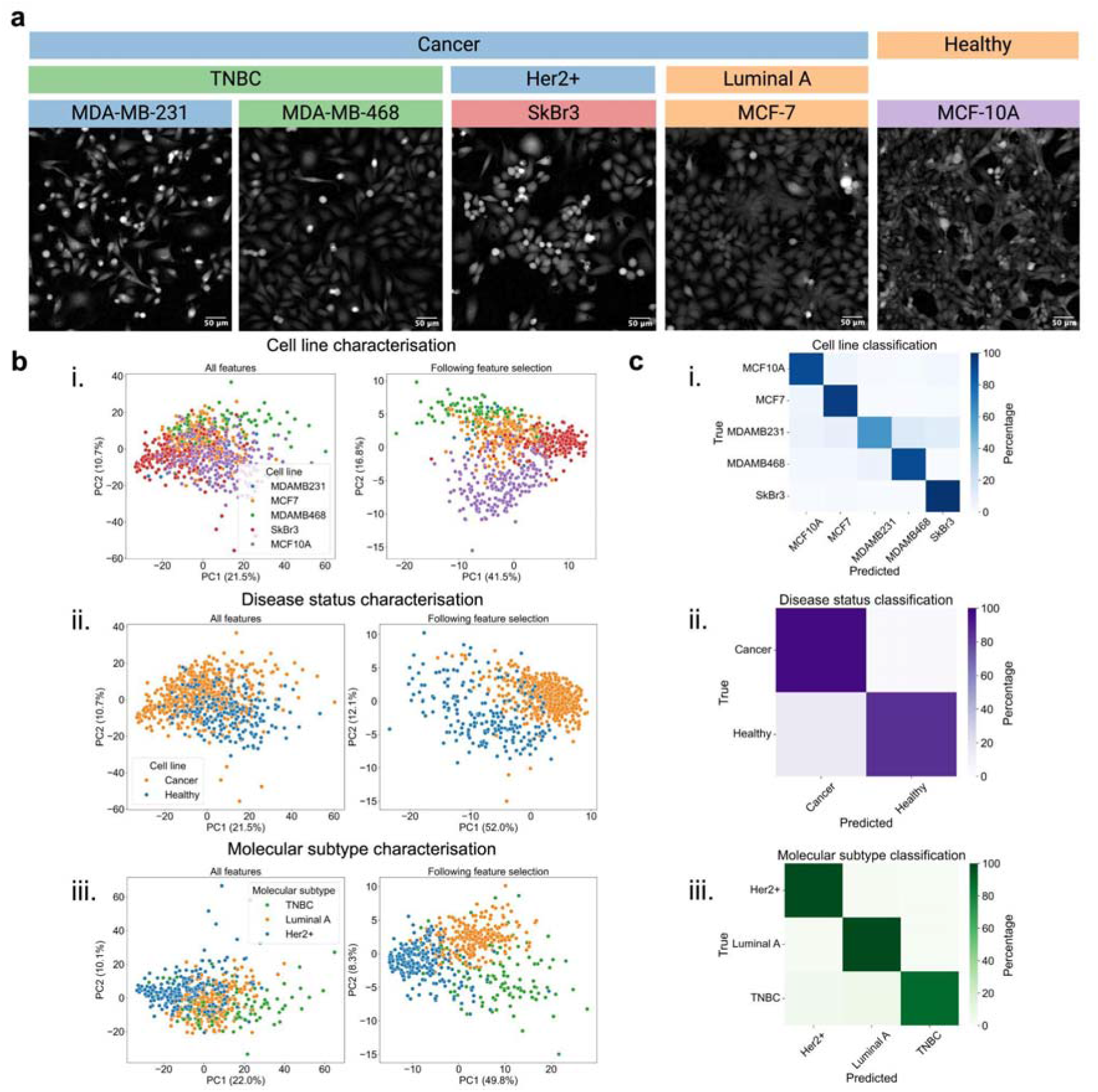
An example of multiclass classification through CellPhePy. **a**. Snapshots taken from ptychographic timelapse videos of MDA-MB-231, MDA-MB-468, SkBr3, MCF-7 and MCF-10A monocultures. Labels used for the analysis included in this figure are provided in the image headings, coloured as they appear in the PCA scores plots in b. **b**. PCA scores plots before and after feature selection for three characterisation applications: i. cell line characterisation, ii. disease status characterisation and iii. molecular subtype characterisation, demonstrating greater separation of cell populations following custom feature selection. **c**. Confusion matrices of test set classification accuracy percentages for XGBoost classification of i. cell lines, ii. disease status, and iii. molecular subtype.

Separation scores are recalculated to capture molecular subtype distinctions, yielding an optimal separation threshold of 0.28 and 141 features achieving separation scores above this threshold. **Figure 3b**(iii) demonstrates the success of this approach in label-free characterisation of molecular subtypes based on phenotypic data.

CellPhePy’s classification capabilities were applied across three different analytical aims: cell line classification, disease status classification, and molecular subtype classification. For each of these objectives, an XGBoost model was trained using the relevant features, and the model was then used to classify unseen data (**Figure 3c**). For this analysis, classification accuracy percentages are reported as the TPR ie. the percentage of cells within a class correctly classified as that class. For cell line classification, accuracy percentages were as follows: MCF-10A = 90%, MCF-7 = 94%, MDA-MB-231 = 61%, MDA-MB-468 = 90%, SkBr3 = 98% (sample sizes are detailed in the Methods). For disease status classification, accuracy scores were Cancer = 97%, Healthy = 85%. For molecular subtype classification, accuracy scores were Her2+ = 98%, Luminal A = 98%, TNBC = 89%. This demonstrates how CellPhePy’s flexible classification approach can be easily adapted to different analytical objectives, making it suitable for a range of applications, including diagnosis, drug screening, and biomarker discovery.

### A CellPhePy GUI for enhanced user accessibility

We developed a CellPhePy GUI to make the toolkit accessible to all users, regardless of their programming expertise. The user is required to provide the path to a folder containing a stack of microscopy time-lapse images on the landing page. Following this, they can click a button to process the images which will perform CellPose segmentation, TrackMate tracking, extraction of single-cell features, and CellPhe’s time series summarisation. The resulting feature tables can then be used as input for the “Single Population” and “Multiple Populations” tabs, which provide dashboards for analysing individual populations or comparing multiple populations, respectively (**Figure 4**). The GUI is run locally, giving users full control over their images without the need to upload them anywhere. It is cross-platform and operates within a web browser, ensuring accessibility on different operating systems. The program is openly available onh GitHub (https://github.com/uoy-research/CellPhe-dashboard) and can be run directly using python, or using Docker.

**Figure 4.**
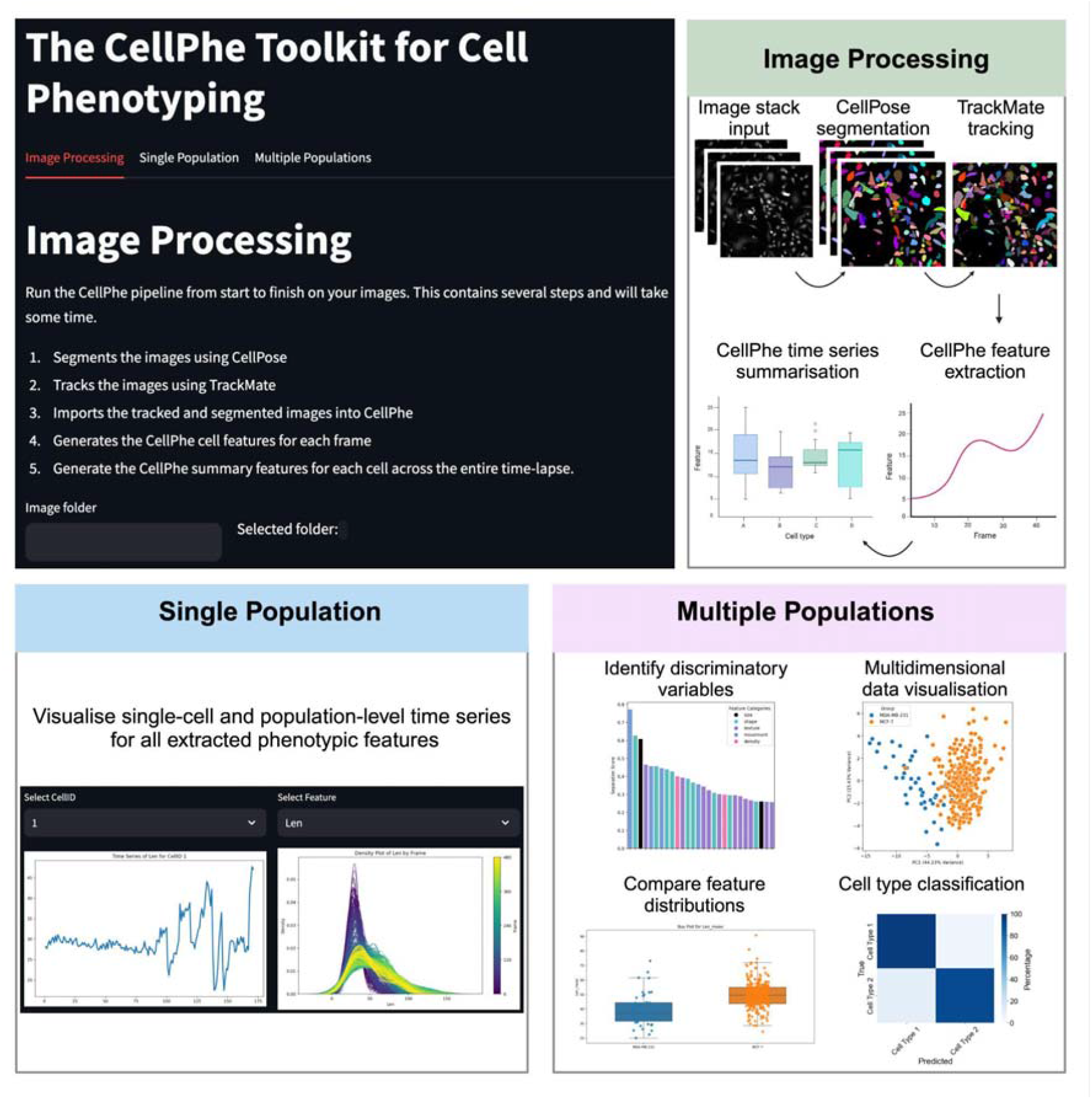
The CellPhePy GUI for enhanced user accessibility. An overview of the three tabs within the CellPhePy GUI: Image Processing, Single Population and Multiple Populations. The image processing tab requires the user to specify a folder path to their stack of microscopy time-lapse images, then automates CellPose segmentation, TrackMate tracking and CellPhe feature extraction. The Single Population tab can be used to visualise single-cell feature time series as well as temporal population-level feature changes. The Multiple Populations tab enables multi-class characterisation and classification of cell types, allowing users to visualise which features are important in distinguishing between their cell populations.

## Discussion

The findings of “2020 BioImage Analysis Survey: community experiences and needs for the future”^13^ highlighted the bioimage analysis community’s desire for developers to improve tool accessibility through comprehensive documentation, intuitive user interfaces, and more user-friendly plugins or packages. In addition, developers were encouraged to make their tools interoperable with popular existing platforms. These recommendations motivated the refactoring of CellPhe to CellPhePy, with the transition from R to Python enabling the tool to integrate with existing python-based tools such as CellPose. Furthermore, incorporation of PyImageJ^14^ enabled headless automation of TrackMate tracking, eliminating the previous requirement to handle this externally through the ImageJ GUI.

Increased modularity of CellPhePy ensures that users are not restricted to a single method for cell segmentation and tracking. For example, users are now able to provide their own labelled masks generated through a software of their choice. Our headless TrackMate tracking code also supports all TrackMate tracking algorithms, including SimpleSparseLAP, SparseLAP, Kalman, AdvancedKalman, NearestNeighbor and Overlap, allowing users to tailor analysis workflows for their specific samples and research needs. This flexibility makes CellPhePy a versatile analysis tool for a wide range of imaging modalities, including ptychography, brightfield and fluorescence. Users can simply replace CellPhePy’s default use of CellPose’s Cyto model with another built-in CellPose model listed on the CellPose documentation page, or use an alternative segmentation method more appropriate for their sample and imaging modality. Users can also add their own custom features within CellPhePy’s feature extraction script, enabling temporal analysis of more sample- and modality-specific features such as changes in intensity or co-localisation of fluorescence markers, for example.

Extending multiclass cell type characterisation and classification capabilities within CellPhePy significantly broadens its potential applications, enabling analysis within more complex experimental setups like large-scale drug screening assays or co-culture experiments. Data can also be repurposed by redefining classes to handle different research questions. For instance, as demonstrated here, groups can be redefined to reveal phenotypic differences between cancerous and healthy states, as well as to distinguish molecular subtypes, thereby increasing its utility for disease diagnosis or study of heterogeneity. We emphasise that multiclass classification may not always be the most suitable approach for a given research question, and this will be reflected in suboptimal results. For example, here we demonstrated that classifying all breast cell lines individually led to poor classification accuracy for MDA-MB-231 cells, with a proportion being misclassified as MDA-MB-468 cells due to shared characteristics between these TNBC cell lines. In this case, a more effective approach would be to use a secondary classifier that focuses on identifying differences between MDA-MB-231 and MDA-MB-468, specifically highlighting the phenotypic distinctions between these two cell lines.

In conclusion, herein we have provided a description of the new python CellPhePy implementation, providing a more adaptable, easier to use version of software for wider uptake of time-lapse imaging data analysis across modalities.

### Materials and Methods. Cell culture

The MDA-MB-231 cells were a gift from M. Djamgoz, Imperial College, London. SkBr3 cells were a gift from J. Rae, University of Michigan and MCF-10A cells were a gift from N. Maitland, University of York. MDA-MB-468 and MCF-7 cells were from ATCC. The molecular identity of all cell lines was verified by short tandem repeat analysis^15^. Authenticated cell stocks were stored in liquid nitrogen and thawed for use in experiments. Thawed cells were subcultured 4-5 times prior to discarding and thawing a new stock to ensure that the molecular identity of cells was retained throughout. All cell lines, except for MCF-10A, were cultured separately in Dulbecco’s Modified Eagle Medium (DMEM) supplemented with 5% fetal bovine serum (FBS) and 4 mM L-glutamine. MCF-10A cells were cultured in DMEM/F12 (Invitrogen) with 5% heat-inactivated horse serum, 0.5 μg/mL hydrocortisone, 20 ng/mL human EGF, 10 μg/mL insulin, and 100 ng/mL cholera toxin. To minimise imaging artefacts, FBS was filtered using a 0.22 μm syringe filter before use. Cells were incubated at 37°C in plastic filter-cap T-25 flasks and were split at a 1:6 ratio during passaging. No antibiotics were added to the cell culture medium. Cells were confirmed to be free of mycoplasma before use in experiments through routine DAPI testing at monthly intervals.

### Image acquisition and exportation

On the day of imaging, cells were placed onto the Phasefocus Livecyte 2 (Phasefocus Limited, Sheffield, UK) and incubated for 30 minutes before image acquisition to allow for temperature equilibration. A 500 μm x 500 μm field of view per well was imaged to capture as many cells, and therefore data observations, as possible. Selected wells were imaged in parallel for ∼22 hours at 20x magnification with 6-minute intervals between frames, resulting in full time-lapses of 222 frames per imaged well. For phase images, Phasefocus’ Cell Analysis Toolbox® software was utilised for image processing, such as use of the rolling ball algorithm to reduce background noise, as well as image exportation.

### Cell type classification

For the multiclass classification examples presented in this study, the training data comprised 43 MDA-MB-231, 73 MDA-MB-468, 213 SkBr3, 205 MCF-10A, and 268 MCF-7 cells. The corresponding test data included 36 MDA-MB-231, 48 MDA-MB-468, 161 SkBr3, 205 MCF-10A, and 294 MCF-7 cells. Classification models were trained, and test sets were evaluated using CellPhePy’s `classify_cells()` function. A confusion matrix summarising classification accuracy percentages was generated using the `sklearn.metrics.confusion_matrix()` function.

Train and test data were divided into different groupings to facilitate three analytical objectives: cell line classification, disease status classification, and molecular subtype classification. In these examples, the data were split as follows:

- Cell line classification: MDA-MB-231, MDA-MB-468, SkBr3, MCF-10A, and MCF-7.
- Disease status classification: Cancer (MDA-MB-231, MDA-MB-468, SkBr3, MCF-7) vs. Healthy (MCF-10A).
- Molecular subtype classification: TNBC (MDA-MB-231, MDA-MB-468) vs. Her2+ (SkBr3) vs. Luminal A (MCF-7).

### CellPhePy

#### Multiclass separation scores

Let *G* be the number of classes. *n*_*g*_ denotes the sample size of class *g, <u>x</u>*_*g*_ denotes the sample mean of a feature for class *g* and 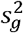 denotes the sample variance of a feature for class *g*.

The grand mean of a feature is given by:

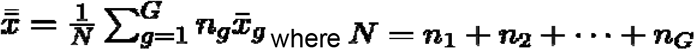

The feature’s between-class variance is given by:

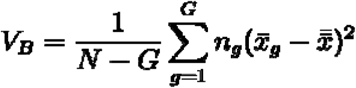

And the feature’s within-class variance is given by:

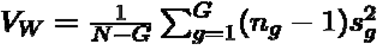

The feature’s separation score can then be calculated by:

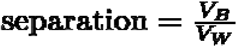

#### Determining optimal separation score threshold

To determine the optimal subset of features, we applied an elbow method based on feature separation scores. Separation scores for all features were sorted in descending order, and a straight line was interpolated between the maximum and minimum scores. For each feature, the vertical distance between its separation score and the interpolated line was calculated. The “elbow point” was identified as the feature with the maximum distance from the line, representing the transition from steep score improvement to a plateau. We then define the optimal separation score threshold as this elbow point, with all downstream classification tasks including only features with separation scores greater than or equal to the optimal threshold.

#### CellPose segmentation

CellPose integration is provided through the official python package: cellpose (https://pypi.org/project/cellpose/, version 3.1.0)

#### TrackMate tracking

Object tracking is provided by the TrackMate ImageJ plugin, interfaced via the PyImagej python package (https://pypi.org/project/pyimagej/, version 1.5.0). This package leverages the scyjava python package (https://pypi.org/project/scyjava/, version 1.10.0) to interface between the Java Virtual Machine running ImageJ, and Python code.

#### The CellPhePy GUI

The CellPhePy GUI is developed using the Streamlit framework, available as a python package (https://pypi.org/project/streamlit/, version 1.41.0). The GUI can be run natively as a Python script, or as part of a Docker image that bundles all the dependencies.

## Author contributions

**Conceptualisation:** LW, WB, P O’T, BC, SL, **Data Curation:** LW, GP, JM, SL, **Formal Analysis**: LW, **Funding Acquisition**: LW, WB, P O’T, BC, BP, **Software:** LW, SL, BC, **Supervision**: LW, WB, P O’T, **Validation:** LW, SL, **Visualisation**: LW, **Writing - Original Draft**: LW, **Writing - Review and Editing**: LW, WB, SL, BC, BP

## Availability of data and materials

The CellPhePy python package is available to install through PyPi at https://pypi.org/project/cellphe/ or through GitHub at https://github.com/uoy-research/CellPhePy. The CellPhePy GUI is also hosted on GitHub (https://github.com/uoy-research/CellPhe-dashboard) and can be installed through the terminal or Docker.

